# On a wing and a prayer: limitations and gaps in global bat wing morphology trait data

**DOI:** 10.1101/2020.12.07.414276

**Authors:** Matt Crane, Inês Silva, Matthew J. Grainger, George A. Gale

## Abstract

Species’ life history traits have a wide variety of applications in ecological and conservation research, particularly when assessing threats. The development and growth of global species trait databases are critical for improving trait-based analyses; however, it is vital to understand the gaps and biases of available data. Bats are an extremely diverse and widely distributed mammalian order, with many species facing local declines and extinction. We conducted a literature review for bat wing morphology, specifically wing loading and aspect ratio, to identify issues with data reporting and ambiguity. We collected data on field methodology, trait terminology, and data reporting and quality. We found several issues regarding semantic ambiguity in trait definitions and data reporting. Globally we found that bat wing morphology trait coverage was low. Only six bat families had over 40% trait coverage, and of those none consisted of more than 11 total species. We found similar biases in trait coverage across IUCN Redlist categories with threatened species having lower coverage. Geographically, North America, Europe, and the Indomalayan regions showed higher overall trait coverage, while both the Afrotropical and Neotropical ecoregions showed poor trait coverage. The underlying biases and gaps with bat wing morphology data have implications for researchers conducting global trait-based assessments. Implementing imputation techniques may address missing data, but only for smaller regional subsets with substantial trait coverage. However, due to the low overall trait coverage, increasing species representation in the database should be prioritized. We suggest adopting an Ecological Trait Standard vocabulary to reduce semantic ambiguity in bat wing morphology traits to improve data compilation and clarity. Additionally, we advocate that researchers adopt an Open Science approach to facilitate the growth of a bat wing morphology trait database.

## 1. Introduction

Bats are an incredibly diverse group constituting nearly one-fifth of all mammals and the second most diverse group of mammals globally. Bats provide a wide range of ecosystem services such as seed dispersal (Melo et al., 2009; Seltzer, Ndangalasi & Cordeiro, 2013) and pest control (Kunz et al, 2011; Maas et al, 2016). Globally, bat populations are under threat from a wide range of pressures (O’Shea et al., 2016; Frick, Kingston & Flanders, 2020) such as habitat loss and degradation (Meyer et al., 2016), wind farms (Arnett el al, 2016), and hunting (Mickleburgh, Waylen & Racey, 2009). Despite their ecological importance and conservation threats, bats have a notable lack of information when compared to other mammals and birds (Frick, Kingston & Flanders, 2020).

Species traits, here defined as measurable characteristics of an organism, provide a wide range of applications for understanding systematics and macroevolution (Harmon et al., 2010; Zamudio, Bell & Mason, 2016) as well as species conservation (Jones, Purvis & Gittleman, 2003; Pacifici et al., 2015). Traits provide an important link to the functional role of species within an ecosystem, and as such trait-based models for vulnerability are a valuable tool (Pacifici et al., 2015; Ramirez-Bautista et al., 2020). However, the ability to apply trait-based assessments relies on having the appropriate trait coverage. Hortal et al (2015) presented the concept of the Raunkiaeran shortfall to describe the lack of knowledge on ecologically relevant species traits, as an important factor to consider when applying large-scale ecological analyses. Biases in the underlying trait data availability can lead to subsequent biases in any analyses. Typically, vertebrate trait data availability is biased taxonomically towards large more widely distributed species and with less representation for reptiles and amphibians, and geographically with less representation in biologically diverse tropical regions (González-Suárez, Lucas & Revilla, 2012; Etard, Morrill & Newbold, 2020). Understanding the underlying biases in a species trait or collection of traits is critical in assessing their applicability to any analysis.

Wing morphology is an important trait in bats, as it links many different natural history traits such as diet (Norberg & Rayner, 1987; Norberg & Fenton, 1988), habitat association (Brigham, Francis & Hamdorf, 1997; Jennings et al., 2004), and range size (Luo et al., 2019). Bat wing morphology is traditionally described using aspect ratio and wing loading (Norberg & Rayner, 1987). For bats wing morphology, aspect ratio is typically defined as the wingspan squared divided by wing area, which describes flight efficiency and maneuverability (Norberg & Rayner, 1987). Higher aspect ratios (e.g. short, wide wings), correspond with faster turning but less efficient flight compared with higher aspect ratios (e.g. long, narrow wings) seen in more efficient flight such as soaring (Norberg & Rayner, 1987; Norberg, Brooke & Trewhella, 2000). Wing loading is calculated by dividing the weight (mass * gravity) of the bat by the wing area and provides a measure of lift potential and influences maneuverability and flight speed. In addition to identifying natural history traits, bat wing morphology has been used for conservation to predict extinct risk (Jones, Purvis & Gittleman, 2003), habitat fragmentation vulnerability (Farneda et al, 2015), and association with urbanization (Jung and Threlfall, 2018). Thus, wing morphology is likely a useful metric for any trait-based assessment with bats.

Globally available databases for species traits are proliferating, as they provide an open repository for assessment (Jones et al., 2009; Kattge et al, 2011; Oliveira et al., 2017). Norberg and Raynor (1987), compiled a global wing morphology database for bats and since then other studies have attempted to compile bat wing morphology either for specific regions (Marinello & Bernard, 2014) or globally (Jones, Purvis & Gittleman, 2003); however, a single centralized and open database does not currently exist for bat wing morphology. Prior to compiling a unified trait database, several key factors must be considered. First, compiled data should use standardized measurement and recording protocols among studies, including following a well-defined and standardized vocabulary (Schneider et al., 2019; Gallagher et al, 2020). Additionally, global trait databases typically contain large volumes of missing data. Missing data can occur both randomly and non-randomly (e.g. geographically, taxonomically) creating potential biases that need to be understood to appropriately validate any results when using a data set, however, missing data can also be imputed to generate larger usable data sets (Taugourdeau et al., 2014).

Our paper seeks to analyze issues associated with developing a global bat wing morphology database. Through a comprehensive literature review, we identify reporting and methodological issues for comparability. Understanding the underlying biases in global trait data is essential in assessing that ability to apply it to global analyses as well as highlight important regions or species to prioritize for study. We created a synthesized dataset for bat wing morphological traits, and then assessed underlying biases in species data availability.

## 2. Methodology

### 2.1 Literature review

In September 2019 and again in February 2020 we searched Scopus and Web of Science databases for articles relating to bat wing morphology, Additionally, we searched Google Scholar using Publish or Perish (Harzing, 2007) software keeping only the first 300 results (Haddaway et al., 2015). For each database we used the following terms: “wing” AND (“morphology” OR “shape” OR “loading” or “measurements”) AND (“bat*” or “chiroptera”). Finally, we also included studies found through reference lists in papers identified from the systematic searches.

To screen papers, we adopted the PRISMA reporting procedure outlined in by Moher, Telzlaff, and Altman (2009). We excluded any papers that did not 1) study bat wing morphology (e.g. birds or robotics), 2) use wing morphology in their study (e.g. only reference wing morphology in the Discussion) or 3) collect primary wing morphology (e.g. reviews and meta-analysis).

From each paper included in the analysis, we collected data on the basic methodology including whether the study used live bats or specimens, the wing area measurement method (e.g. photo, tracing), and whether they used only primary data. Norberg & Raynor (1987) noted important semantic ambiguity in the term “wing area”, and whether the body and/or tail membrane is included. We followed Bininda-Emonds & Russell, 1994 and use two distinct terms lifting surface area (LSA) and wing area (WA). Lifting surface area is defined following “wing area” set out by Norberg & Raynor (1987) which includes the area of the tail membrane, body excluding the head, and wing. We define wing area specifically referring to the area of the patagium only. To assess methods reporting quality we recorded whether the paper explicitly reported whether they used LSA or WA. When the paper was unclear, we attempted to extrapolate which definition was used based on provided figures or references.

We also collected information on data reporting and whether the study provided data for both direct field measures, mass, forearm, wingspan, and wing area as well as calculated variables, aspect ratio, wing loading, and relative wing loading. We considered data that were only displayed in a figure as absent because raw values could not be extracted. Recent work supports importance of incorporating and providing intraspecific trait variability (Guralnick et al., 2016; Kissling et al., 2018), thus we also evaluated the reported data quality based on whether the study provided raw data for individuals, means and standard deviations, or only as means without SDs or sample sizes.

### 2.2 Data availability and biases

We assessed global data availability for bat wing morphology to assess taxonomic and geographic trends. First, we validated and updated the species names from the literature as older papers often used outdated scientific names; however, as different taxonomic backbones exist, we retained the original verbatim scientific name provided. As we investigated patterns in conservation status based on the IUCN Redlist, we corrected our scientific names to match the IUCN convention. We first checked for synonyms in the GBIF backbone naming system, and then followed the IUCN naming convention to allow for extracting threat status. We downloaded data on geographic realm and IUCN Redlist Conservation Status for each species, using package “rredlist” (Chamberlain, 2020). We did not include data when individuals were not identified to species in the study, nor species that did not occur on the IUCN species list nor the Catalogue of Life. For valid species that did not occur on the IUCN Redlist we manually added the conservation status (Data Deficient) and realm (following the IUCN convention). We followed the terminology presented in Etard et al. (2020) for assessing trait data availability, with trait coverage referring to the proportion of missing data for a specific trait, and trait completeness which is the proportion of traits have an available estimate for a given species. We then evaluated trait coverage and completeness based on conservation status, taxonomic family, and realm to review patterns.

We assessed spatial patterns in trait coverage by first descriptively comparing the overall trait coverage for the ecoregions Palearctic, Nearctic, Neotropical, Afrotropical, Indomalayan, and Australasian ecoregions (following the WWF Ecoregions classification; Olson et al., 2004). Since the Palearctic realm covers a large geographic area, we further divided the area by geopolitical divisions into Asiatic Palearctic (Asiatic Russia and Eastern Asian), European Palearctic, Middle Eastern Palearctic (consisting of Central and Southwestern Asia), and Palearctic Africa. Additionally, we reviewed the assemblage-level trait coverage for wing loading and aspect ratio for each ecoregion. We overlapped species distributions from IUCN (Spatial Range Dataset IUCN, 2020; downloaded October, 2020) and calculated the assemblage-level trait coverage globally using a grid size of approximately 50 km by 50 km using the package “fasterize” in R (Ross, 2020). We then extracted the median and interquartile range for trait coverage values in each ecoregion.

### 2.3 Trait data extraction, validation, and compilation

When available, we extracted morphological data to compile a unified data set. As our objective was to compile a generalized database on bat species wing morphology, we first combined data for species within studies (e.g. when studies reported values for different sexes). For studies that used wing area instead of LSA we used the correction based on taxonomic family used by Norberg and Raynor (1987) which added the tail and body area as a percentage of reported wing area (20% Emballonurids, Rhinolophids, Hipposiderids, and Mormoopids, 25% for Verpertilionids, and 16% for Phyllostomidids).

When wing loading and/or relative wing loading was not reported but mass and wing area were, we estimated a mean wing loading and considered the estimate as a single observation, and thus did not calculate standard deviation. We also applied this procedure to estimate aspect ratio and relative wing loading. We estimated wing loading from relative wing loading and vice versa when only mass and either wing loading or relative wing loading were reported.

We created a compiled data set with wing morphology data based on the reviewed literature. Prior to combining estimates for species from different studies we identified outlier records and either corrected or removed the data. To identify outliers, we first calculated the mean standard deviations for aspect ratio and wing loading. We defined an outlier as any species that exhibit a standard deviation above 1.5 times the mean. For the species we identified we reviewed all data from studies reporting for those species to determine whether the study exhibited systematic biases. For studies that exhibiting systematic biases we sought to identify and then rectify the issue, and if we could not then the data from the study was excluded from the final global data set. For individual species outliers (e.g. when the study did not exhibit systematic biases) we removed records that did not report sample size and standard deviation. We did not remove outliers, if the variation in the data arose potentially due to geographic or taxonomic variation (e.g. different sub-species). Additionally, when forearm and/or were missing for a species, we used values from the PanTHERIA database (Jones et al., 2009) when available. We used the finalized data set to evaluate patterns in missing data taxonomically, geographically, and in conservation status.

### 2.4 Methodological impact assessment

We investigated the effect of area measurement method and area definition on wing area/lift surface area values. From our methodological review, we only included studies that clearly reported how wing area was defined and how wing area was measured. Next, we only included species that had area measurements from at least two different methods and specifically stated in the methods that they used the lifting surface area definition for “wing area”. The available data did not provide a large enough sample to make viable comparisons between wing area definitions for different families (only 5 species from 3 genera had measurements for both definitions). We ran Bayesian generalized linear mixed model using the package “*brms*” in R Studio (Bürkner, 2017), with a lognormal likelihood, and 6 chains with 6,000 iterations each (1,000 set as burn-in). We kept default flat priors, with measurement method (tracing vs. photograph) set as a fix8282.ed effect, and species and study ID as random effects. We checked for model convergence using trace plots and R-hat values, and assessed model fit through posterior predictive checks, leave-one-out cross-validation and Bayesian R^2^ (Bürkner, 2017). We ran all analyses and visualizations in R Studio using R v3.6.3 (R Core Team, 2020)

## 3. Results

### 3.1 Method variation and reproducible reporting

Our search returned 738 references and yielded 147 sources after screening and full-text assessment (Supp File S1) covering 508 species from 18 families. Area measurement techniques varied, but most studies either used wing tracings (n = 71) or photographs (n = 49), with ~10% of studies not reporting how area measurements were obtained (n = 15; Figure 1). Other methods included using photos of bats in flight, or directly estimating area in the field with a planimeter. Roughly half of the studies reviewed clearly distinguished whether they used LSA or wing area (56.4%), while 24.8% did not report methods in a way that an area definition could be extrapolated (Figure 1). All but three studies used the term “wing area”, regardless of the area definition described in the methodology.

**Figure 1.**
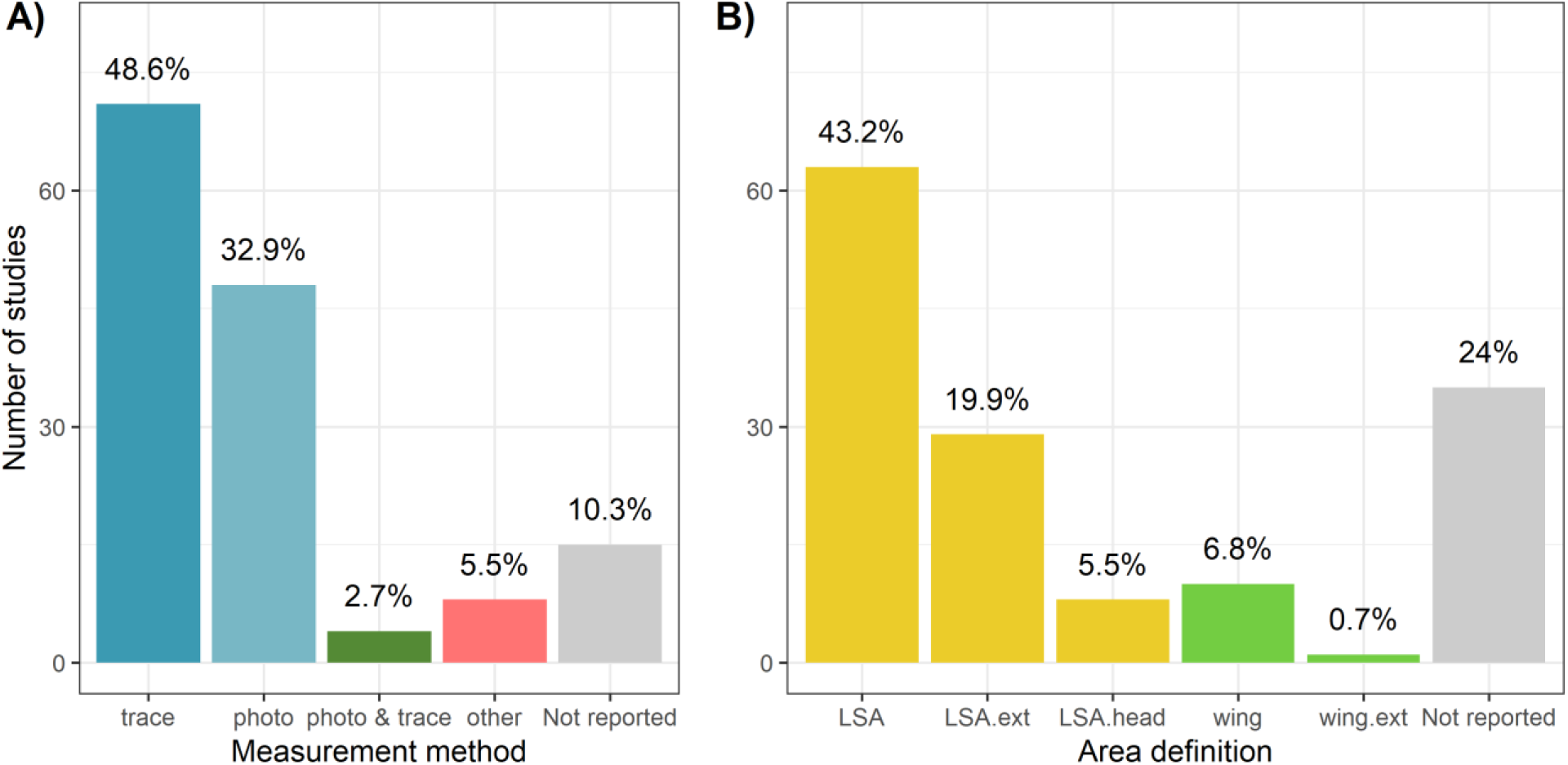
Methods used to measure wing area in bats (A), and how studies defined wing area either using lift surface area (LSA) or wing area (wing) with “ext” denoting that the definition was extrapolated based on the provided reference. Lift surface area includes the area of the patagium uropatagium, and the body excluding the head, and wing area refers to only the area of the patagium.

Overall, 83 studies (56.5 %) provided all the relevant trait measurements (mass, wingspan, wing area) for calculating wing loading, relative wing loading, and aspect ratio. Conversely, 19 studies did not report data for directly measured traits, and a further five studies did not report any raw data despite using the data for analysis. For directly measured traits, both wingspan and wing area exhibited higher missing data rates than mass (Figure 2).

**Figure 2.**
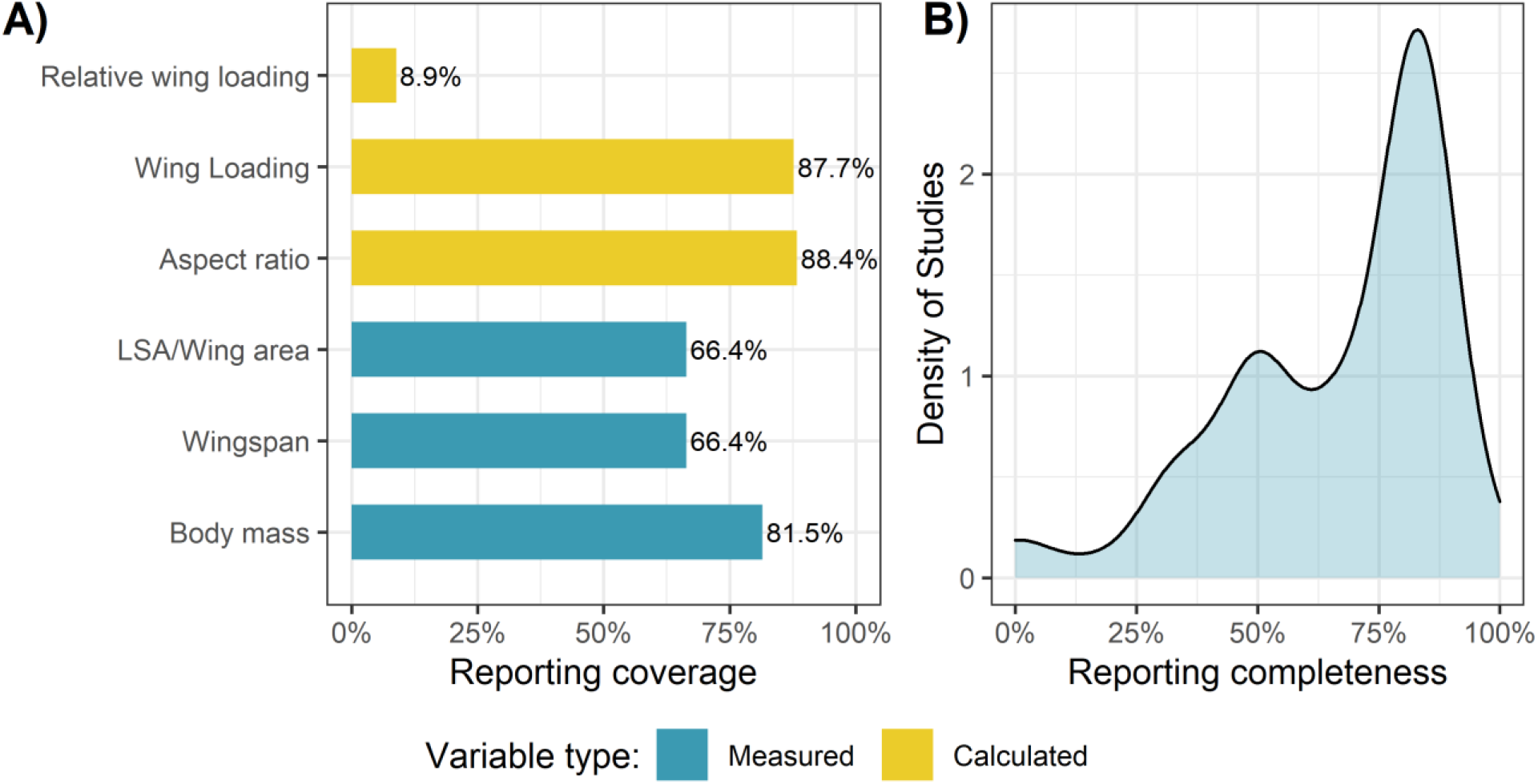
A) Reporting coverage for each bat wing morphology trait and the B) density of reporting completeness for studies that collected bat wing morphology traits.

Measured traits had a higher missing rate than calculated variables, except for relative wing loading, which was not typically calculated (only 15 of 147 studies calculated RWL with 13 reporting values). Of the 13 studies that reported relative wing loading, two studies did not provide the base measurements mass and/or wing area to estimate wing loading. However, the data loss of those two studies resulted in 66 records and 14 unique species not having available wing loading data. Similarly, 19 studies that did not report relative wing loading also did not provide the necessary data to calculate relative wing loading, leading to 24 unique species without relative wing loading estimates. Not reporting mass led to a loss of 38 species with complete wing morphology data from the final database. From the literature we identified 487 species which had estimates for both wing loading and aspect ratio. Overall, reporting issues and methodological inconsistencies led to 141 of the studied species (~28%) to have no wing loading nor aspect ratio data in the final data set.

When provided, data quality also varied with only 14 of 147 studies providing complete individual level data; however, of those studies only two provided the data as an available supplement. Most studies (71.4%) reported species-level mean and a measure of dispersion (either standard deviation or standard error), and a smaller portion only reporting a mean (9.5%).

### 3.2 Influence of field measurement method on area

From the available data, 47 species had records for lift surface area using both tracing and photographs allowing for direct comparison. The models converged, showing caterpillar-like trace plots, and all R-hat values approximated 1. Posterior predictive checks indicated that predicted and observed data had similar distributions, and reflected adequate model fit. However, Pareto shape k parameters indicated the presence of outliers with high influence for both models (Supp. File S2). Measuring lift surface area using a tracing compared to photographs did not show any effect on area estimate (Table 1; Supp. File S2).

**Table 1.** Bayesian regression outputs for the model comparing between measuring lift surface area using a tracing or a photograph.

| Model structure | Estimate | Est.Error | L95%<br>CI | U95%<br>CI | Rhat | Bulk<br>ESS | Tail<br>ESS |
| --- | --- | --- | --- | --- | --- | --- | --- |
| *Group-Level Effects:* |  |  |  |  |  |  |  |
| sd(Reference ID) | 0.09 | 0.03 | 0.04 | 0.16 | 1 | 4602 | 7051 |
| sd(Scientific Name) | 0.49 | 0.05 | 0.40 | 0.61 | 1 | 2803 | 6403 |
| *Population-Level Effects:* |  |  |  |  |  |  |  |
| Intercept | 4.96 | 0.08 | 4.80 | 5.12 | 1 | 1960 | 4458 |
| method MeasurementTrace | 0.01 | 0.06 | -0.09 | 0.13 | 1 | 7001 | 10561 |
| Bayes R<sup>2</sup> |  |  |  |  |  | 94.99% |  |

From the collected wing morphology data, only 6 species (*Cynopterus sphinx*, *Cynopterus brachyotis*, *Cynopterus horsefieldii*, *Molossus molossus*, and *Pteropus tonganus*) had records using both definitions of area. The primary difference between lift surface area and wing area is the inclusion of the uropatagium in LSA and from the available species only *Molossus molossus* has a relatively large uropatagium. As such, while modeling was not possible to determine the differences between using wing area or lift surface area on wing loading and aspect ratio, we did see a difference in wing loading (LSA definition mean wing loading = 19.7 Nm^−2^; wing area definition mean wing loading = 16.5) and aspect ratio (LSA aspect ratio = 7.7; wing area aspect ratio = 8.7) estimates for *Molossus molossus*. The observed pattern is consistent theoretically considering that as the tail area increases in relation to the size of the wing area both aspect ratio and wing loading will decrease as they are inversely proportional to area.

### 3.3 Global species data availability

After removing outlier records and correcting for methodology we compiled a dataset comprised of 430 species with at least some available wing morphology data (wingspan, wing area, wing loading, relative wing loading or aspect ratio), representing approximately 27.8 % of global bat species. When data was available, trait completeness was high as 311 of the 430 species had estimates for all relevant traits (mass, wingspan, wing area, aspect ratio, wing loading, and relative wing loading). However, overall wing morphology data showed low trait coverage across most bat families, with only 7 families showing ≥ 50% data availability and only one of those families (Mormoopidae) consisted of more than 6 species (Figure 3). For genus level trait coverage, 79 genera (36.1%) had no species with available data. Within families that had at least one species with available data, the percent of genera without any data ranged from 22.6% to 66.7% (Suppl Table S1). Primarily, species wing loading data were derived from one or two studies (78.7%), while only 23 species had five or more references. Aspect ratio showed a similar pattern with only 25 species with five or more references.

**Figure 3.**
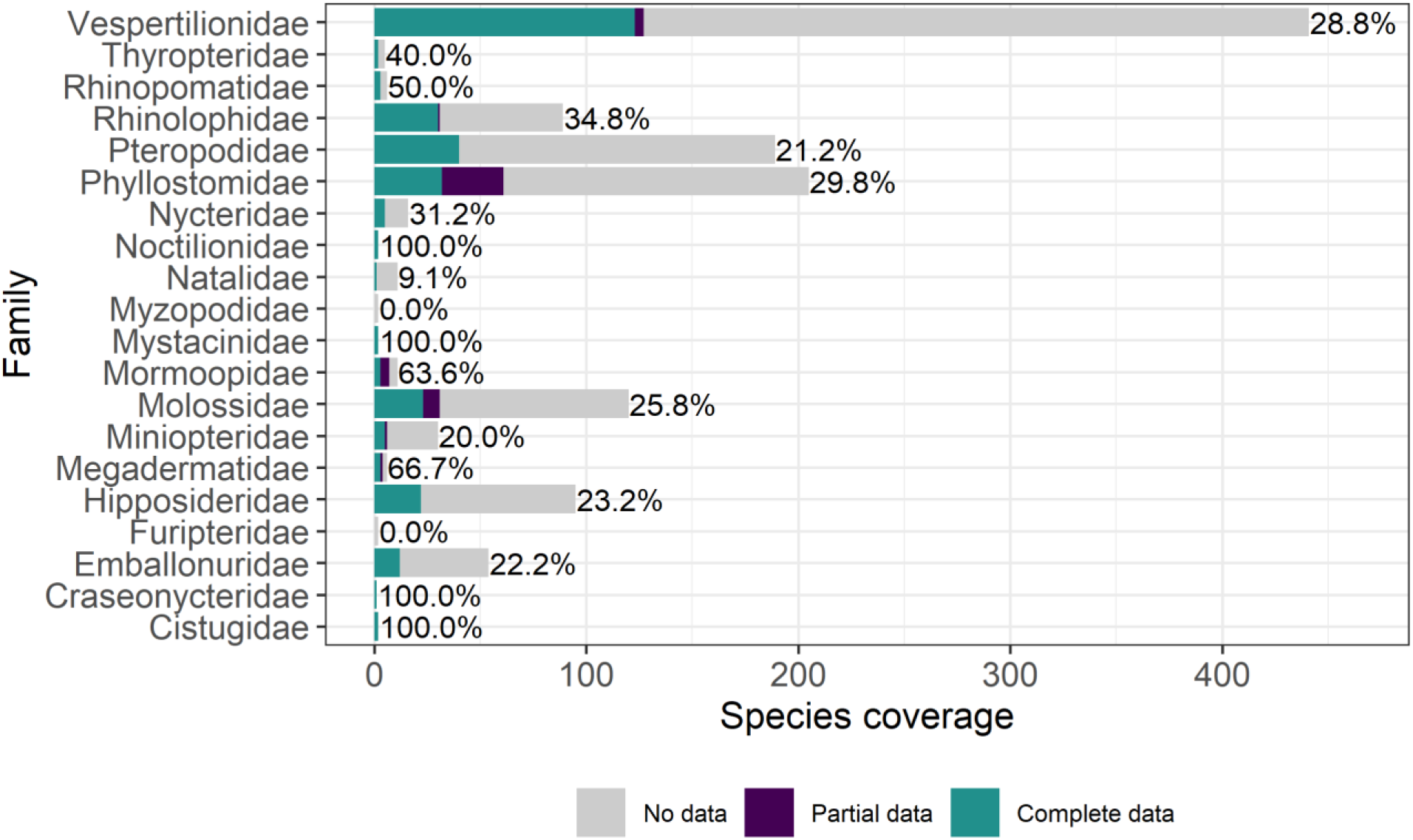
Wing morphology data availability for bat species by family, showing species with no data (e.g. without wing loading and aspect ratio), partial data (wing loading and aspect ratio but not mass, wingspan, and/or lift surface area) or complete data (all relevant trait data available) along with the percent of species within each family that have at least some data available.

Across all IUCN categories wing morphology data exhibited a high volume of missingness (Figure 4). The distribution of available data follows a similar pattern to overall species categorization. Only 17 of the 193 species listed in a Threatened category (Vulnerable, Endangered, Critically Endangered), have wing morphology data and all 17 had full trait completeness (i.e. available data for mass, wingspan, wing area, wing loading, relative wing loading, and aspect ratio). When looking at listed threats under the IUCN assessment for threatened species, we saw similar patterns of low trait coverage across threat categories. Additionally, all the threats identified as high impact from the IUCN assessment had no wing morphology data available (Supp. Table S2).

**Figure 4.**
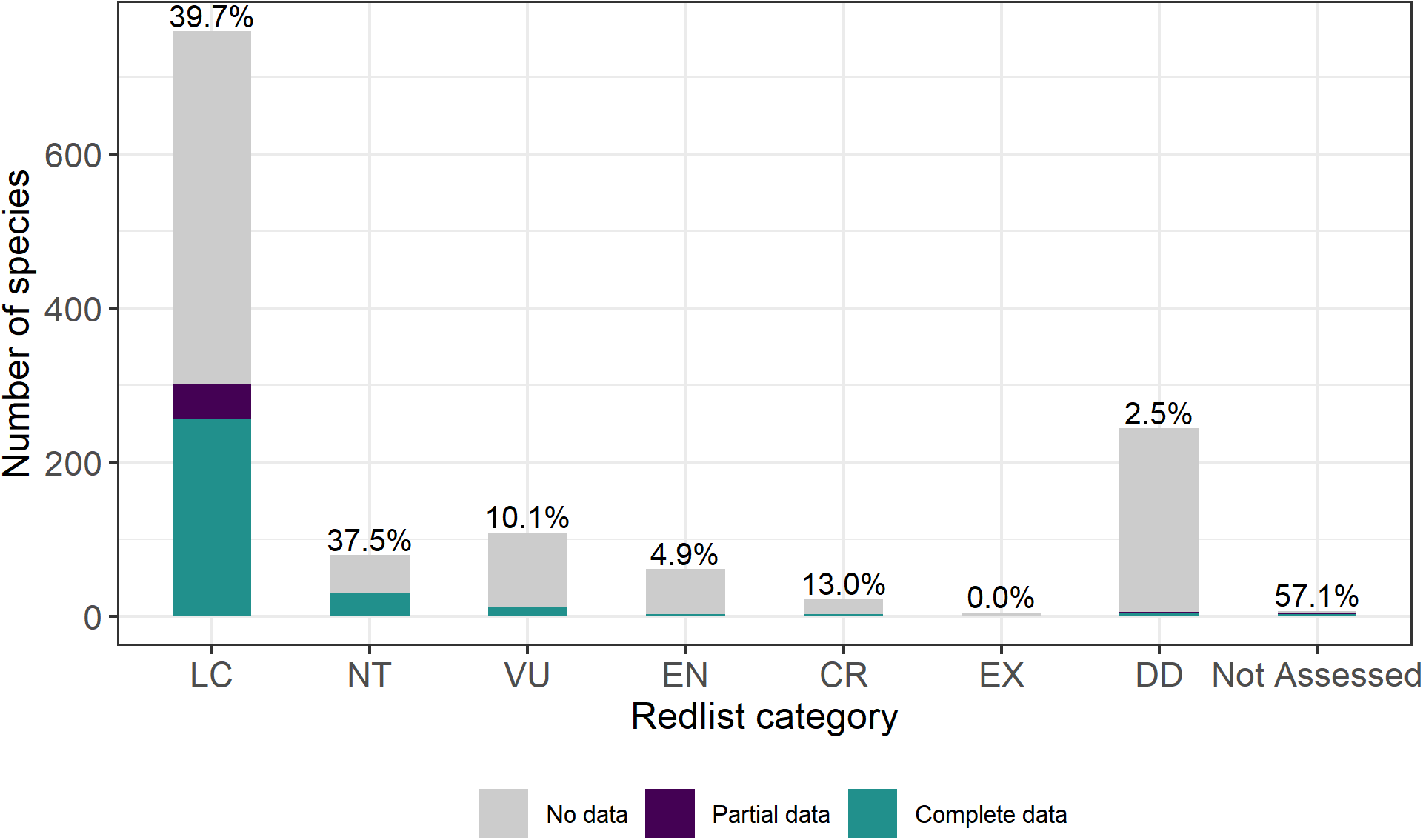
Bat wing morphology data availability (species with both wing loading and aspect ratio) for IUCN Redlist category Least Concern (LC), Near Threatened (NT), Vulnerable (VU), Endangered (EN), Critically Endangered (CR), Extinct (EX), Data Deficient (DD), and species not currently assessed by the IUCN. Percentages above the bars indicates percent of species with at least some available trait data for each category.

Trait coverage for wing loading and aspect ratio varied between ecoregions but was typically low overall (Supplemental Table S3). Only the Nearctic realm exhibited trait coverage over 50% with an ecoregion wide trait coverage of 65.5% for both wing morphology and aspect ratio. The Nearctic also exhibited the highest median assemblage-level trait coverage for both wing loading and aspect ratio, followed by the Palearctic and Indomalayan realms; however, several realms showed high variation in assemblage-level trait coverage (Figure 5b). Trait coverage varied within the geopolitical regions we defined for the Palearctic, with the Asiatic Palearctic exhibiting lower median trait coverage than the other regions (Figure 5c). The global wing morphology dataset showed geographic biases with North America and Europe having higher overall trait coverage for species, while Asiatic Palearctic, tropical Australasia, and tropical Africa had relatively lower data coverage.

**Figure 5.**
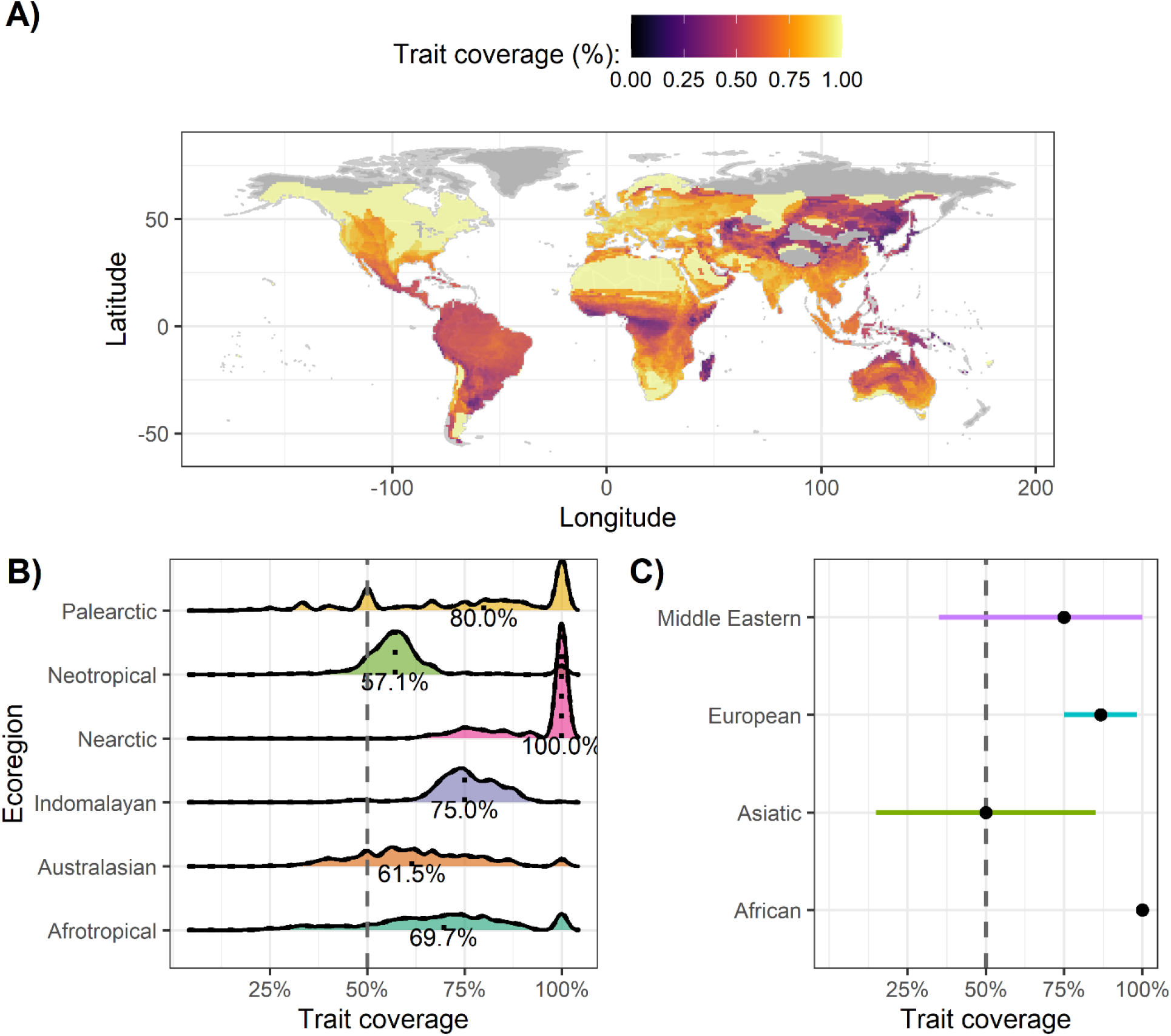
(A) Global assemblage-level wing loading trait coverage. Darker colors highlight regions with relatively fewer number of species with available data. Light grey indicates that either no bat species are present or that range data was not available. (B) Assemblage-level wing loading trait coverage density for each with the median coverage value. (C) Median and interquartile range of wing loading trait coverage for geopolitical regions within the Palearctic ecoregion.

## 4. Discussion

Overall, bat wing morphology data exhibits poor trait coverage, with a small proportion (< 30%) of species with available data for each trait we assessed. We identified several underlying issues within the wing morphology literature that inhibit consolidating a global database. From the available data, bat wing morphology trait coverage showed both taxonomic and geographic biases which limits assessing conservation status and threat risks globally with the currently available data.

While we found variation in field techniques for measuring wing area, most studies adopted one of two methods which produced similar area estimates. One of the biggest issues arose from semantic ambiguity in terminology, primarily wing area definition, and this should be addressed. We suggest following Schneider et al., (2019) and adopting an Ecological Trait-data Standard Vocabulary (ETS), which as a minimum requires studies to report a value (column named *traitValue*), standard units for the value (*traitUnit*), a unified descriptive name (*traitName*), and a scientific taxon (*scientificName*). Using the standard column naming is not required but facilitates faster inclusion into a unified database. Trait name is important for maintaining consistency across studies, and for bat wing morphology we propose a set of unambiguous trait names (Table 2; Figure 6).

**Table 2.** Proposed unified terminology for bat wing morphology traits, with the current potentially ambiguous terms the suggested unified trait name, the suggested trait unit, and an unambiguous definition for the proposed trait

| Current terms | unified traitName | traitUnit | Description |
| --- | --- | --- | --- |
| Wingspan | wingspan | cm | The linear distance from wing tip to wing tip or measured as the distance from the midline of the body to a single wing tip doubled |
| Wing area; Lifting surface area | wing_area | cm<sup>2</sup> | The full area of the patagium, directly measured from a wing tracing or photograph |
|  | lifting_surface_area | cm<sup>2</sup> | The total area of the patagium, uropatagium (if present), and body excluding the head |
|  | wing_area_rec | cm<sup>2</sup> | The rectangular approximation of the patagium based on the wing length (measured from the edge of the body to the wing tip) multiplied by the wing wing width (measured starting at the fifth digit down to the tip of the fifth digit) |
| Aspect ratio | partial_aspect_ratio | NA | The wingspan squared divided by wing_area |
|  | total_aspect_ratio | NA | The wingspan squared divided by lifting_surface_area |
| Wing loading | partial_wing_loading | Nm<sup>-2</sup> | mass x gravitational constant divided by wing_area |
|  | total_wing_loading | Nm<sup>-2</sup> | mass x gravitational constant divided by lifting_surface_area |
| Relative wing loading; geometric wing loading | relative_partial_loading | NA | partial_wing_loading divided by mass raised to 1/3. Note that mass and not weight (mass * g) should be used |
|  | relative_total_loading | NA | total_wing_loading divided by mass raised to 1/3. Note that mass and not weight (mass * g) should be used |

**Figure 6.**
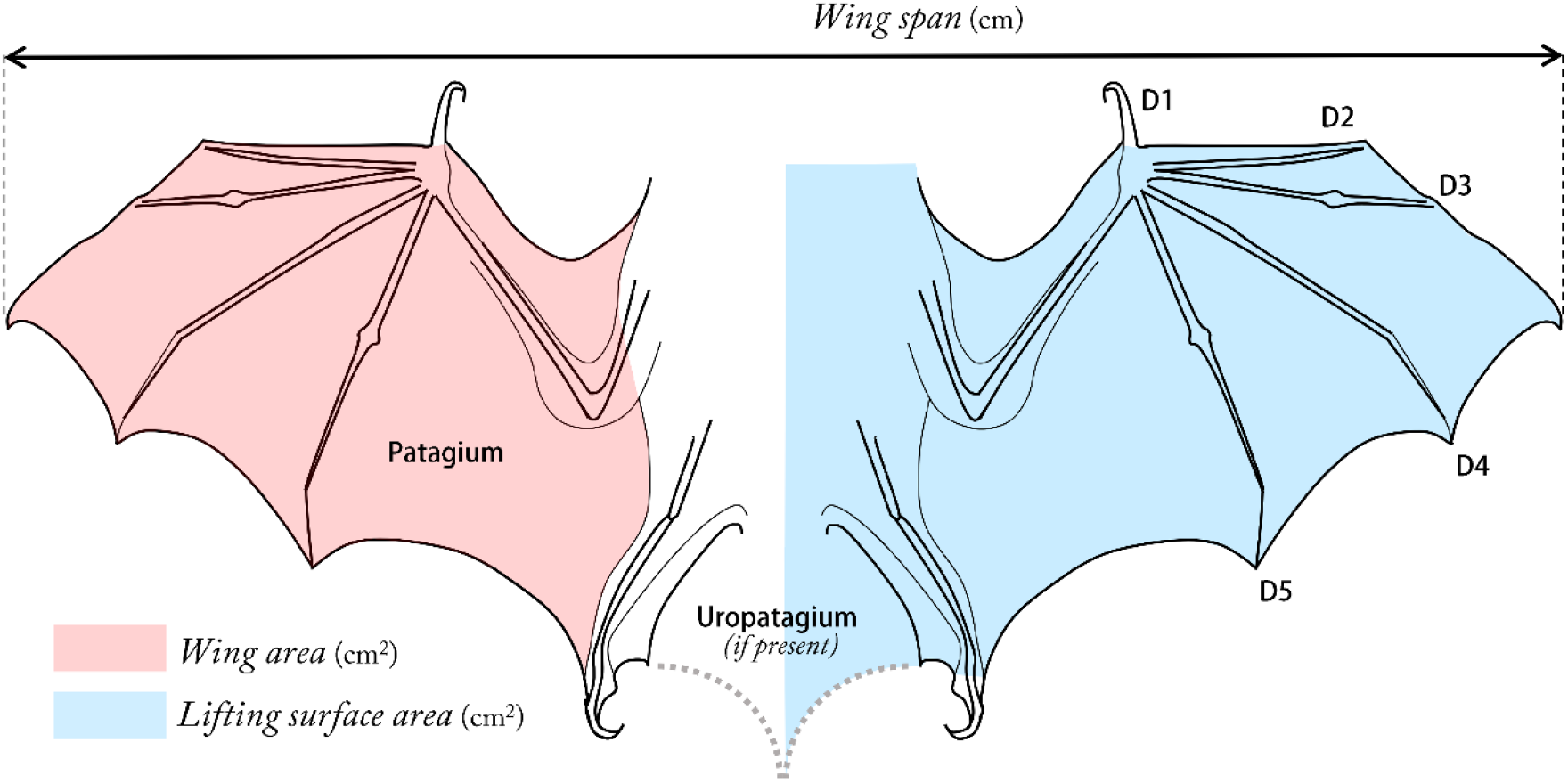
Bat wing outline designating regions included in the "wing area" definition (left) and lifting surface area (right).

We also identified numerous gaps in data reporting across studies. We documented that primary morphological traits such as forearm, mass, wingspan, and wing area were underreported in the literature. Additionally, few studies (14) provided fully open available individual level data, and more problematically several studies (13) either provided no data, did not provide species level data, or only presented data in a figure. Open Science can facilitate the development and accelerate the growth of trait datasets (Gallagher et al 2020), thus we suggest that studies collecting bat wing morphology traits should adopt an Open Science approach to address the data reporting gap. Omitting trait values limits the ability to unify trait data and estimate additional variables. For instance, we saw that not reporting mass along with either wing loading or relative wing loading meant that we were not able to generate an estimate for both traits. Additionally, while we focused on wing loading and aspect ratio, several other methods exist for comparing bat wing morphology such as geometric analysis (Dietz, Dietz & Siemers, 2006; Schmieder et al., 2015), and indices based on skeletal measures (Findley, Studier & Wilson, 1972; Bader et al., 2015). Providing values for all relevant traits measured in a study, can help either estimate or even calculate other trait values strengthening the overall data availability.

Wing morphology data are not available for most bat species across multiple families and regions. Geographically, several regions showed low data availability particularly South America, Madagascar, parts of Oceania, the Middle East, and Asiatic Palearctic. The geographic biases raise several concerns with applying a global approach to meta-analysis using wing morphology data. Bat populations face a myriad of threats which vary depending on the region (Frick, Kingston & Flanders, 2020). We found a similar trend in low trait coverage for threatened species (González-Suárez, Lucas & Revilla, 2012). We identified gaps in data coverage for IUCN threatened category species, as well as when considering trait coverage for specific high impact threats. Given the underlying low trait coverage and biases, it is critical for researchers using bat wing morphology to consider the missingness mechanism within their framework (Nakagawa & Freckleton, 2008; Baraldi & Enders, 2010). For instance, missing data patterns within conservation status and specific threats could confound results and inhibit conservation-based conclusions

Missing data pattens (e.g. whether data is missing at random or not) within available trait data, also relates to how ecologists should handle missing data cases (Nakagawa & Freckleton, 2008). Missing data rarely occurs completely at random, which means that simple approaches such as case deletion can introduce strong biases (Nakagawa & Freckleton, 2008; González-Suárez, Lucas & Revilla, 2012; Pakeman, 2014). As missing data is ubiquitous across fields, multiple new techniques have been developed to impute non-random missing data (Bruggeman, Heringa, & Brandt, 2009; Pantanowitz & Marwala, 2009; van Buuren & Groothuis-Oudshoorn, 2010). A major concern with applying any imputation with is the volume of missing data relative to available data. Penone et al. (2014) showed that some techniques could handle up to 40% missing data when traits were phylogenetically conserved. In addition to accounting for phylogeny, relationships between traits can be used in imputation models. Incorporating additional traits to impute bat wing morphology data may provide a promising approach, as other traits are globally well sampled for mammals; however, it unclear whether this is the case specifically for bats (Etard et al., 2020). Globally trait coverage for bat wing morphology does not meet the 40% criteria, but studies focused on a smaller subset could still benefit from data imputation.

### Caveats

We did not conduct our literature review searches in additional languages, which may change the data availability coverage especially in regions where English is not the primary language. Some of the geographic biases we observed may be a factor of limiting our search language to English. English is not the primary language in several of the regions with low trait coverage. However, our English search provides a large starting centralized dataset and additionally provides a framework for adding bat wing morphology data from non-English sources and from grey literature. Additionally, the English search did return three papers in Chinese. We advocate for using the data presented in this study only as a starting point for developing a more robust global repository for bat wing morphology. As the global database grows it will be important to revisit our assessment of taxonomic and geographic biases. To this end we have provided R code to easily facilitate future reevaluations, as well as a bat wing morphology database with three levels: individual level data, study level data, and pooled species level data.

## Supporting information

Supplemental File 2_Taxonomic trait coverage table

## Acknowledgements

We would like the thank King Mongkut’s Petchra Pra Jom Klao Scholarship for supporting the project.

## Notes

http://doi.org/10.5281/zenodo.4309198

### Competing Interest Statement

The authors have declared no competing interest.

## References

Arnett EB, Baerwald EF, Mathews F, Rodrigues L, Rodríguez-Durán A, Rydell J, Villegas-Patraca R, Voigt CC. 2016. Impacts of wind energy development on bats: a global perspective. In: Bats in the Anthropocene: conservation of bats in a changing world. Springer Cham, 295–323.

Bader E, Jung K, Kalko EKV, Page RA, Rodriguez R, Sattler T. 2015. Mobility explains the response of aerial insectivorous bats to anthropogenic habitat change in the Neotropics. Biological Conservation 186:97–106. DOI: 10.1016/j.biocon.2015.02.028.

Baraldi AN, Enders CK. 2010. An introduction to modern missing data analyses. Journal of School Psychology 48:5–37. DOI: 10.1016/j.jsp.2009.10.001.

Bininda-Emonds ORP, Russell AP. 1994. Flight style in bats as predicted from wing morphometry: the effects of specimen preservation. Journal of Zoology 234:275–287. DOI: 10.1111/j.1469-7998.1994.tb06075.x.

Brigham RM, Francis RL, Hamdorf S. 1997. Microhabitat Use by Two Species of Nyctophilus Bats: a Test of Ecomorphology Theory. Australian Journal of Zoology 45:553. DOI: 10.1071/ZO97026.

Bruggeman J, Heringa J, Brandt BW. 2009. PhyloPars: estimation of missing parameter values using phylogeny. Nucleic Acids Research 37:W179–W184. DOI: 10.1093/nar/gkp370.

Bürkner P-C. 2017. brms: An R package for Bayesian multilevel models using Stan. Journal of statistical software 80:1–28.

Chamberlain S. (2020). rredlist: ‘IUCN’ Red List Client. R package version 0.6.0. https://CRAN.R-project.org/package=rredlist

Dietz C, Dietz I, Siemers BM. 2006. Wing Measurement Variations In The Five European Horseshoe Bat Species (Chiroptera: Rhinolophidae). Journal of Mammalogy 87:1241–1251. DOI: 10.1644/05-MAMM-A-299R2.1.

Etard A, Morrill S, Newbold T. 2020. Global gaps in trait data for terrestrial vertebrates. Global Ecology and Biogeography:geb.13184. DOI: 10.1111/geb.13184.

Farneda FZ, Rocha R, López-Baucells A, Groenenberg M, Silva I, Palmeirim JM, Bobrowiec PED, Meyer CFJ. 2015. Trait-related responses to habitat fragmentation in Amazonian bats. Journal of Applied Ecology 52:1381–1391. DOI: 10.1111/1365-2664.12490.

Frick WF, Kingston T, Flanders J. 2020. A review of the major threats and challenges to global bat conservation. Annals of the New York Academy of Sciences 1469:5–25. DOI: 10.1111/nyas.14045.

Gallagher RV, Falster DS, Maitner BS, Salguero-Gómez R, Vandvik V, Pearse WD, Schneider FD, Kattge J, Poelen JH, Madin JS, Ankenbrand MJ, Penone C, Feng X, Adams VM, Alroy J, Andrew SC, Balk MA, Bland LM, Boyle BL, Bravo-Avila CH, Brennan I, Carthey AJR, Catullo R, Cavazos BR, Conde DA, Chown SL, Fadrique B, Gibb H, Halbritter AH, Hammock J, Hogan JA, Holewa H, Hope M, Iversen CM, Jochum M, Kearney M, Keller A, Mabee P, Manning P, McCormack L, Michaletz ST, Park DS, Perez TM, Pineda-Munoz S, Ray CA, Rossetto M, Sauquet H, Sparrow B, Spasojevic MJ, Telford RJ, Tobias JA, Violle C, Walls R, Weiss KCB, Westoby M, Wright IJ, Enquist BJ. 2020. Open Science principles for accelerating trait-based science across the Tree of Life. Nature Ecology & Evolution 4:294–303. DOI: 10.1038/s41559-020-1109-6.

González-Suárez M, Lucas PM, Revilla E. 2012. Biases in comparative analyses of extinction risk: mind the gap: *Data biases in comparative analyses*. Journal of Animal Ecology 81:1211–1222. DOI: 10.1111/j.1365-2656.2012.01999.x.

Guralnick RP, Zermoglio PF, Wieczorek J, LaFrance R, Bloom D, & Russell L. (2016). The importance of digitized biocollections as a source of trait data and a new VertNet resource. Database, 2016 baw158. https://doi.org/10.1093/database/baw158

Haddaway NR, Collins AM, Coughlin D, Kirk S. 2015. The Role of Google Scholar in Evidence Reviews and Its Applicability to Grey Literature Searching. PLOS ONE 10:e0138237. DOI: 10.1371/journal.pone.0138237.

Harmon LJ, Losos JB, Jonathan Davies T, Gillespie RG, Gittleman JL, Bryan Jennings W, Kozak KH, McPeek MA, Moreno-Roark F, Near TJ, Purvis A, Ricklefs RE, Schluter D, Schulte II JA, Seehausen O, Sidlauskas BL, Torres-Carvajal O, Weir JT, Mooers AØ. 2010. Early Bursts Of Body Size And Shape Evolution Are Rare In Comparative Data. Evolution 64:2385–2396. DOI: 10.1111/j.1558-5646.2010.01025.x.

Harzing, AW. 2007. Publish or Perish, available from https://harzing.com/resources/publish-or-perish

Hortal J, de Bello F, Diniz-Filho JAF, Lewinsohn TM, Lobo JM, Ladle RJ. 2015. Seven Shortfalls that Beset Large-Scale Knowledge of Biodiversity. Annual Review of Ecology, Evolution, and Systematics 46:523–549. DOI: 10.1146/annurev-ecolsys-112414-054400.

Jennings NV, Parsons S, Barlow KE, Gannon MR. 2004. Echolocation Calls and Wing Morphology of Bats from the West Indies. Acta Chiropterologica 6:75–90. DOI: 10.3161/001.006.0106.

Jones KE, Bielby J, Cardillo M, Fritz SA, O’Dell J, Orme CDL, Safi K, Sechrest W, Boakes EH, Carbone C, Connolly C, Cutts MJ, Foster JK, Grenyer R, Habib M, Plaster CA, Price SA, Rigby EA, Rist J, Teacher A, Bininda-Emonds ORP, Gittleman JL, Mace GM, Purvis A. 2009. PanTHERIA: a species-level database of life history, ecology, and geography of extant and recently extinct mammals: Ecological Archives E090–184. Ecology 90:2648–2648. DOI: 10.1890/08-1494.1.

Jones KE, Purvis A, Gittleman JL. 2003. Biological Correlates of Extinction Risk in Bats. The American Naturalist 161:601–614. DOI: 10.1086/368289.

Jung K, Threlfall CG. 2018. Trait-dependent tolerance of bats to urbanization: a global meta-analysis. Proceedings of the Royal Society B: Biological Sciences 285:20181222. DOI: 10.1098/rspb.2018.1222.

Kattge J, Díaz S, Lavorel S, Prentice IC, Leadley P, Bönisch G, Garnier E, Westoby M, Reich PB, Wright IJ, Cornelissen JHC, Violle C, Harrison SP, van Bodegom PM, Reichstein M, Enquist BJ, Soudzilovskaia NA, Ackerly DD, Anand M, Atkin O, Bahn M, Baker TR, Baldocchi D, Bekker R, Blanco CC, Blonder B, Bond WJ, Bradstock R, Bunker DE, Casanoves F, Cavender-Bares J, Chambers JQ, Chapin Iii FS, Chave J, Coomes D, Cornwell WK, Craine JM, Dobrin BH, Duarte L, Durka W, Elser J, Esser G, Estiarte M, Fagan WF, Fang J, Fernández-Méndez F, Fidelis A, Finegan B, Flores O, Ford H, Frank D, Freschet GT, Fyllas NM, Gallagher RV, Green WA, Gutierrez AG, Hickler T, Higgins SI, Hodgson JG, Jalili A, Jansen S, Joly CA, Kerkhoff AJ, Kirkup D, Kitajima K, Kleyer M, Klotz S, Knops JMH, Kramer K, Kühn I, Kurokawa H, Laughlin D, Lee TD, Leishman M, Lens F, Lenz T, Lewis SL, Lloyd J, Llusià J, Louault F, Ma S, Mahecha MD, Manning P, Massad T, Medlyn BE, Messier J, Moles AT, Müller SC, Nadrowski K, Naeem S, Niinemets Ü, Nöllert S, Nüske A, Ogaya R, Oleksyn J, Onipchenko VG, Onoda Y, Ordoñez J, Overbeck G, Ozinga WA, Patiño S, Paula S, Pausas JG, Peñuelas J, Phillips OL, Pillar V, Poorter H, Poorter L, Poschlod P, Prinzing A, Proulx R, Rammig A, Reinsch S, Reu B, Sack L, Salgado-Negret B, Sardans J, Shiodera S, Shipley B, Siefert A, Sosinski E, Soussana J-F, Swaine E, Swenson N, Thompson K, Thornton P, Waldram M, Weiher E, White M, White S, Wright SJ, Yguel B, Zaehle S, Zanne AE, Wirth C. 2011. TRY - a global database of plant traits: TRY - A Global Database of Plant Traits. Global Change Biology 17:2905–2935. DOI: 10.1111/j.1365-2486.2011.02451.x.

Kissling WD, Walls R, Bowser A, Jones MO, Kattge J, Agosti D, Amengual J, Basset A, van Bodegom PM, Cornelissen JHC, Denny EG, Deudero S, Egloff W, Elmendorf SC, Alonso García E, Jones KD, Jones OR, Lavorel S, Lear D, Navarro LM, Pawar S, Pirzl R, Rüger N, Sal S, Salguero-Gómez R, Schigel D, Schulz K-S, Skidmore A, Guralnick RP. 2018. Towards global data products of Essential Biodiversity Variables on species traits. Nature Ecology & Evolution 2:1531–1540. DOI: 10.1038/s41559-018-0667-3.

Kunz TH, Braun de Torrez E, Bauer D, Lobova T, Fleming TH. 2011. Ecosystem services provided by bats: Ecosystem services provided by bats. Annals of the New York Academy of Sciences 1223:1–38. DOI: 10.1111/j.1749-6632.2011.06004.x.

Luo B, Santana SE, Pang Y, Wang M, Xiao Y, Feng J. 2019. Wing morphology predicts geographic range size in vespertilionid bats. Scientific Reports 9:4526. DOI: 10.1038/s41598-019-41125-0.

Maas B, Karp DS, Bumrungsri S, Darras K, Gonthier D, Huang JC-C, Lindell CA, Maine JJ, Mestre L, Michel NL, Morrison EB, Perfecto I, Philpott SM, Şekercioğlu ÇH, Silva RM, Taylor PJ, Tscharntke T, Van Bael SA, Whelan CJ, Williams-Guillén K. 2016. Bird and bat predation services in tropical forests and agroforestry landscapes: Ecosystem services provided by tropical birds and bats. Biological Reviews 91:1081–1101. DOI: 10.1111/brv.12211.

Marinello MM, Bernard E. 2014. Wing morphology of Neotropical bats: a quantitative and qualitative analysis with implications for habitat use. Canadian Journal of Zoology 92:141–147. DOI: 10.1139/cjz-2013-0127.

Melo FPL, Rodriguez-Herrera B, Chazdon RL, Medellin RA, Ceballos GG. 2009. Small Tent-Roosting Bats Promote Dispersal of Large-Seeded Plants in a Neotropical Forest. Biotropica 41:737–743. DOI: 10.1111/j.1744-7429.2009.00528.x.

Meyer CFJ, Struebig MJ, Willig MR. 2016. Responses of tropical bats to habitat fragmentation, logging, and deforestation. In: Bats in the Anthropocene: conservation of bats in a changing world. Springer Cham, 63–103.

Mickleburgh S, Waylen K, Racey P. 2009. Bats as bushmeat: a global review. Oryx 43:217. DOI: 10.1017/S0030605308000938.

Nakagawa S, Freckleton RP. 2008. Missing inaction: the dangers of ignoring missing data. Trends in Ecology & Evolution 23:592–596. DOI: 10.1016/j.tree.2008.06.014.

Norberg UML, Brooke AP, Trewhella WJ. 2000. Soaring and Non-soaring bats of the family Pteropodidae (Flying Foxes, Pteropus spp.): Wing morphology and flight performance. The Journal of Experimental Biology 203:651–664.

Norberg UM, Fenton MB. 1988. Carnivorous bats? Biological Journal of the Linnean Society 33:383–394. DOI: 10.1111/j.1095-8312.1988.tb00451.x..

Norberg UM, Rayner J. 1987. Ecological morphology and flight in bats (Mammalia; Chiroptera): wing adaptations, flight performance, foraging strategy and echolocation. Philosophical Transactions of the Royal Society of London. B, Biological Sciences 316:335–427. DOI: 10.1098/rstb.1987.0030.

Oliveira BF, São-Pedro VA, Santos-Barrera G, Penone C, Costa GC. 2017. AmphiBIO, a global database for amphibian ecological traits. Scientific Data 4:170123. DOI: 10.1038/sdata.2017.123.

O’Shea TJ, Cryan PM, Hayman DTS, Plowright RK, Streicker DG. 2016. Multiple mortality events in bats: a global review: Multiple mortality events in bats. Mammal Review 46:175–190. DOI: 10.1111/mam.12064.

Olson DM, Dinerstein E, Wikramanayake ED, Burgess ND, Powell GVN, Underwood EC, D’Amico JA, Itoua I, Strand HE, Morrison JC, Loucks CJ, Allnutt TF, Ricketts TH, Kura Y, Lamoreux JF, Wettengel WW, Hedao P, and Kassem KR. Terrestrial Ecoregions of the World: A New Map of Life on Earth. BioScience 51:933–938.

Pacifici M, Foden WB, Visconti P, Watson JEM, Butchart SHM, Kovacs KM, Scheffers BR, Hole DG, Martin TG, Akçakaya HR, Corlett RT, Huntley B, Bickford D, Carr JA, Hoffmann AA, Midgley GF, Pearce-Kelly P, Pearson RG, Williams SE, Willis SG, Young B, Rondinini C. 2015. Assessing species vulnerability to climate change. Nature Climate Change 5:215–224. DOI: 10.1038/nclimate2448.

Pakeman RJ. 2014. Functional trait metrics are sensitive to the completeness of the species’ trait data? Methods in Ecology and Evolution 5:9–15. DOI: 10.1111/2041-210X.12136.

Pantanowitz A, Marwala T. 2009. Missing Data Imputation Through the Use of the Random Forest Algorithm. In: Yu W, Sanchez EN eds. Advances in Computational Intelligence. Advances in Intelligent and Soft Computing. Berlin, Heidelberg: Springer Berlin Heidelberg, 53–62. DOI: 10.1007/978-3-642-03156-4_6.

Penone C, Davidson AD, Shoemaker KT, Di Marco M, Rondinini C, Brooks TM, Young BE, Graham CH, Costa GC. 2014. Imputation of missing data in life-history trait datasets: which approach performs the best? Methods in Ecology and Evolution 5:961–970. DOI: 10.1111/2041-210X.12232.

Ross N. 2020. fasterize: Fast Polygon to Raster Conversion. R package version 1.0.3. https://CRAN.R-project.org/package=fasterize

R Core Team. 2020. R: A language and environment for statistical computing. R Foundation for Statistical Computing, Vienna, Austria. URL https://www.R-project.org/.

Ramírez‐Bautista A, Thorne JH, Schwartz MW, Williams JN. 2020. Trait‐based climate vulnerability of native rodents in southwestern Mexico. Ecology and Evolution 10:5864–5876. DOI: 10.1002/ece3.6323.

Schmieder DA, Benítez HA, Borissov IM, Fruciano C. 2015. Bat Species Comparisons Based on External Morphology: A Test of Traditional versus Geometric Morphometric Approaches. PLOS ONE 10:e0127043. DOI: 10.1371/journal.pone.0127043.

Schneider FD, Fichtmueller D, Gossner MM, Güntsch A, Jochum M, König‐Ries B, Le Provost G, Manning P, Ostrowski A, Penone C, Simons NK. 2019. Towards an ecological trait‐ data standard. Methods in Ecology and Evolution 10:2006–2019. DOI: 10.1111/2041-210X.13288.

Seltzer CE, Ndangalasi HJ, Cordeiro NJ. 2013. Seed Dispersal in the Dark: Shedding Light on the Role of Fruit Bats in Africa. Biotropica 45:450–456. DOI: 10.1111/btp.12029.

Taugourdeau S, Villerd J, Plantureux S, Huguenin‐Elie O, Amiaud B. 2014. Filling the gap in functional trait databases: use of ecological hypotheses to replace missing data. Ecology and Evolution 4:944–958. DOI: 10.1002/ece3.989.

Van Buuren S, and Groothuis-Oudshoorn K. 2011. Mice: multivariate imputation by chained equations in R. Journal of Statistical Software. 45:1–67.

Zamudio KR, Bell RC, Mason NA. 2016. Phenotypes in phylogeography: Species’ traits, environmental variation, and vertebrate diversification. Proceedings of the National Academy of Sciences 113:8041–8048. DOI: 10.1073/pnas.1602237113.

