## Supplemental File 1_Figures and Tables for "On a wing and a prayer: limitations and gaps in global bat wing morphology trait data"

### Supplement S1: PRISMA flow diagram for literature review

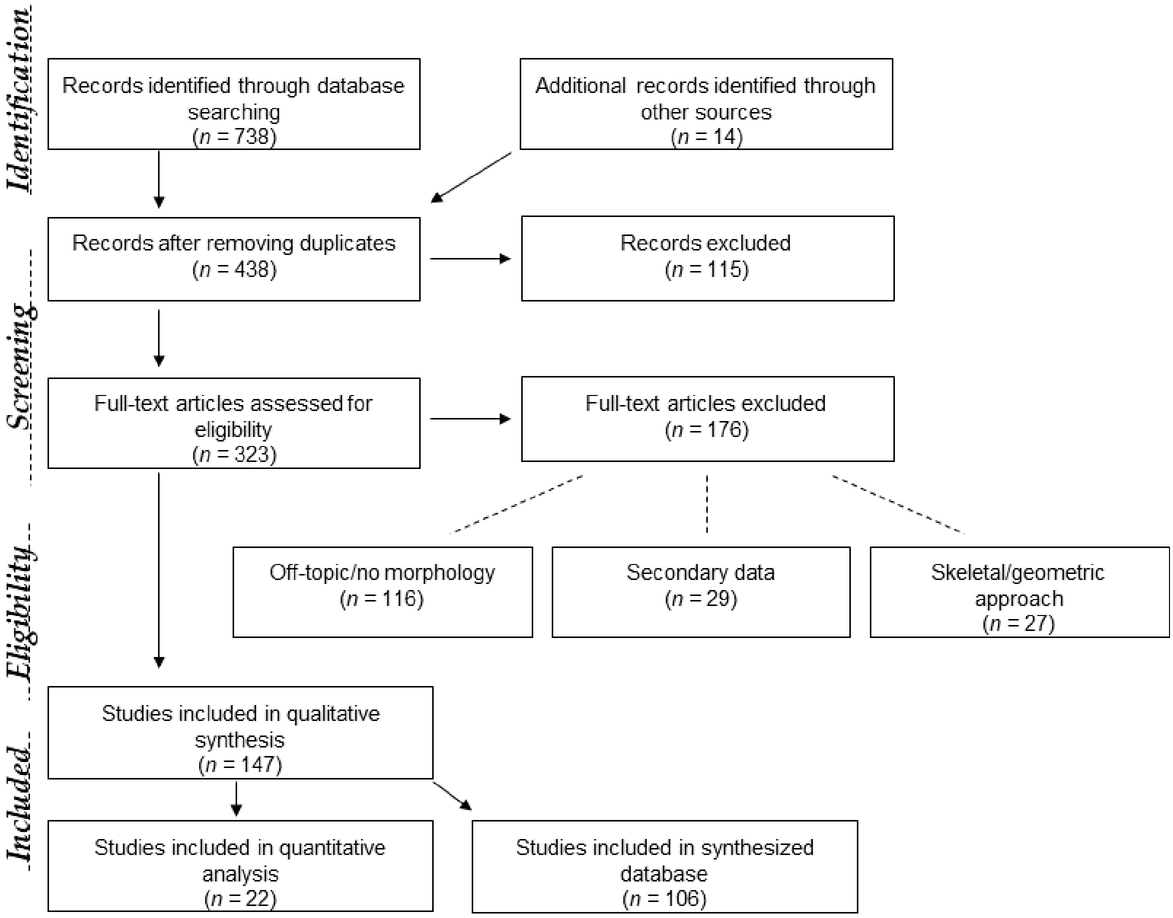


**Supplemental Figure 1** PRISMA flow diagram for structured literature review of bat wing morphology studies. Adopted from: Moher D, Liberati A, Tetzlaff J, Altman DG, The PRISMA Group (2009). Preferred Reporting Items for Systematic Reviews and Meta-Analyses: The PRISMA Statement. PLoS Med 6(7): e1000097. doi:10.1371/journal.pmed1000097

### Supplement S2: Outputs from the Bayesian hierarchical models

Summary output:

```
Family: lognormal
  Links: mu = identity; sigma = identity
Formula: wingArea_cm2 ~ methodMeasurement + (1 | scientificName.iucn) + (1 | refID)
   Data: method.model.data (Number of observations: 139)
Samples: 6 chains, each with iter = 6000; warmup = 1000; thin = 1;
         total post-warmup samples = 30000

Group-Level Effects:
~refID (Number of levels: 21)
              Estimate Est.Error l-95% CI u-95% CI Rhat Bulk_ESS Tail_ESS
sd(Intercept)     0.09      0.03     0.04     0.16 1.00     4602     7051

~scientificName.iucn (Number of levels: 47)
              Estimate Est.Error l-95% CI u-95% CI Rhat Bulk_ESS Tail_ESS
sd(Intercept)     0.49      0.05     0.40     0.61 1.00     2803     6403

Population-Level Effects:
                       Estimate Est.Error l-95% CI u-95% CI Rhat Bulk_ESS Tail_ESS
Intercept                  4.96      0.08     4.80     5.12 1.00     1960     4458
methodMeasurementtrace     0.01      0.06    -0.09     0.13 1.00     7001    10561

Family Specific Parameters:
      Estimate Est.Error l-95% CI u-95% CI Rhat Bulk_ESS Tail_ESS
sigma     0.10      0.01     0.08     0.12 1.00     8214    14224

Samples were drawn using sampling(NUTS). For each parameter, Bulk_ESS
and Tail_ESS are effective sample size measures, and Rhat is the potential
scale reduction factor on split chains (at convergence, Rhat = 1).
```

LOO output:

```
Computed from 30000 by 139 log-likelihood matrix

         Estimate   SE
elpd_loo   -605.1 13.2
p_loo        60.3  7.5
looic      1210.1 26.4
------
Monte Carlo SE of elpd_loo is NA.

Pareto k diagnostic values:
                         Count Pct.    Min. n_eff
(-Inf, 0.5]   (good)     75    54.0%   3631
 (0.5, 0.7]   (ok)       45    32.4%   232
   (0.7, 1]   (bad)      17    12.2%   40
   (1, Inf)   (very bad)  2     1.4%   23
```

Bayes $R^2$:

```
   Estimate   Est.Error      Q2.5     Q97.5
R2 0.949874 0.009887981 0.9268897 0.9650663
```

Trace and Density Plots for MCMC Samples:

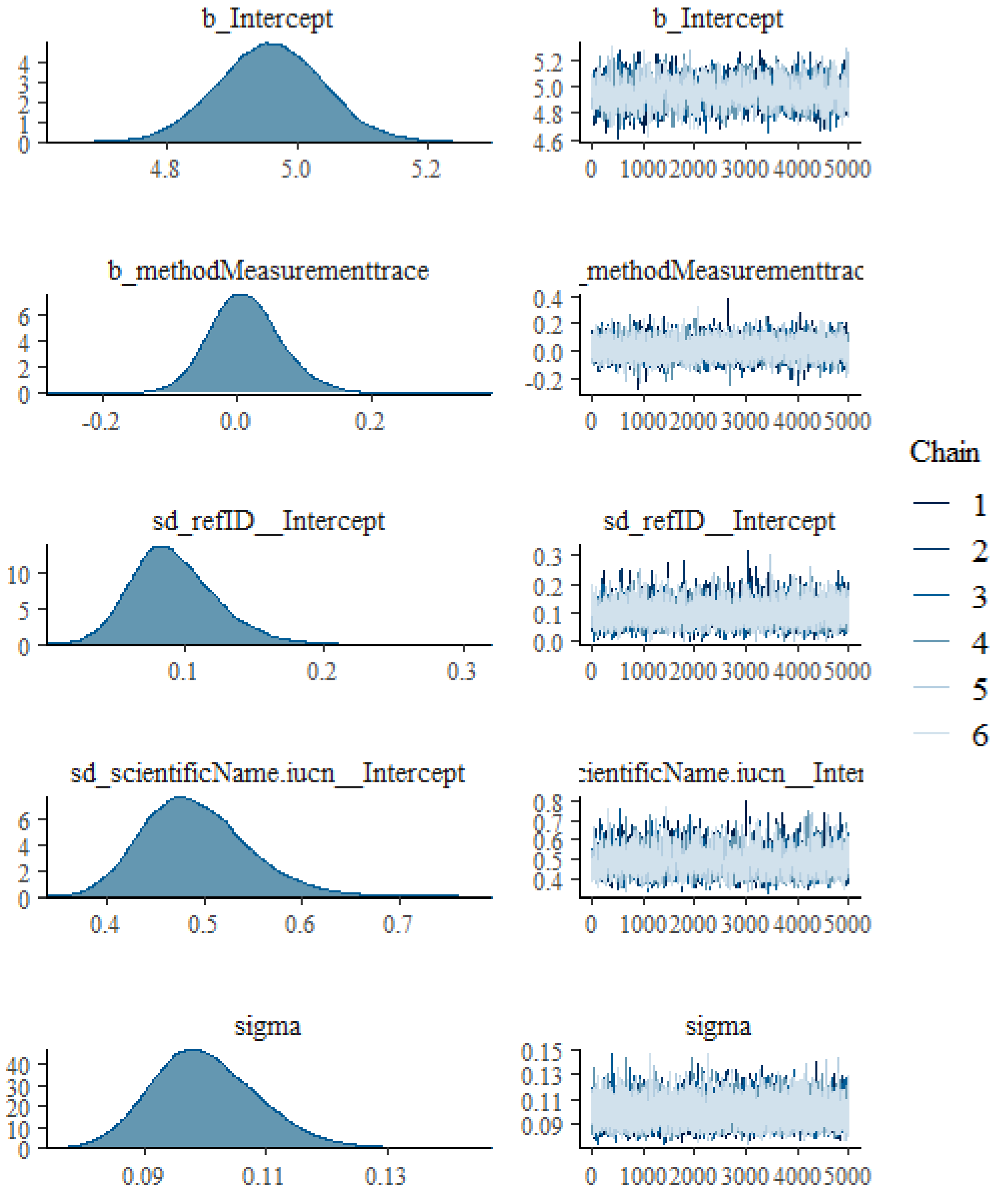


Posterior predictive check:

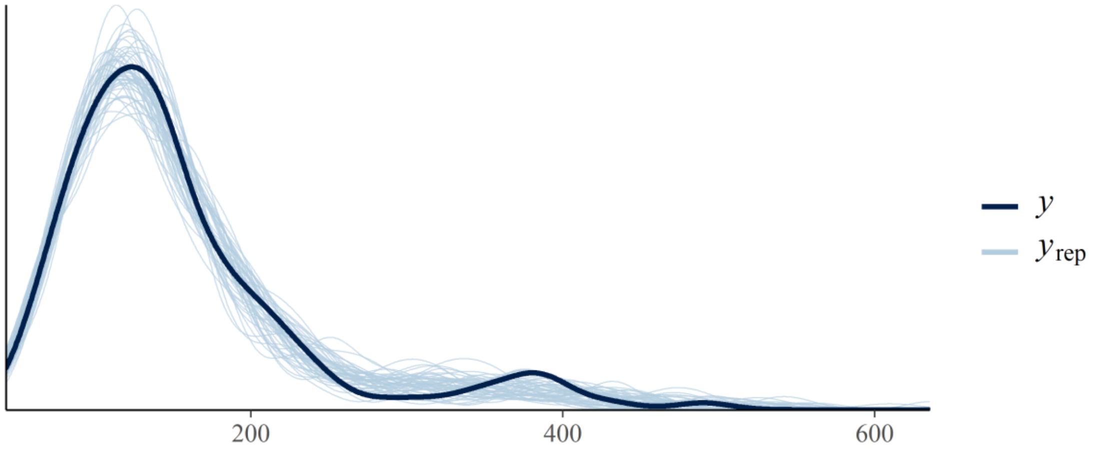

### Supplement S4: Data on IUCN threats and geographic coverage

**Supplemental Table** S2 Trait coverage by IUCN threat impact for aspect ratio, wing loading, and relative wing loading for species listed as Threatened, Endangered, or Critically Endangered under IUCN Redlist

| Threat impact category | Aspect ratio | Wing loading | Relative wing loading |
|---|---|---|---|
| High Impact | 0.0% | 0.0% | 0.0% |
| Low Impact | 11.5% | 13.4% | 13.4% |
| Medium Impact | 5.6% | 5.6% | 5.6% |
| No/Negligible Impact | 0.0% | 0.0% | 0.0% |
| Past Impact | 12.5% | 12.5% | 12.5% |
| Unknown | 2.3% | 2.3% | 2.3% |
| NA | 5.8% | 5.8% | 5.8% |

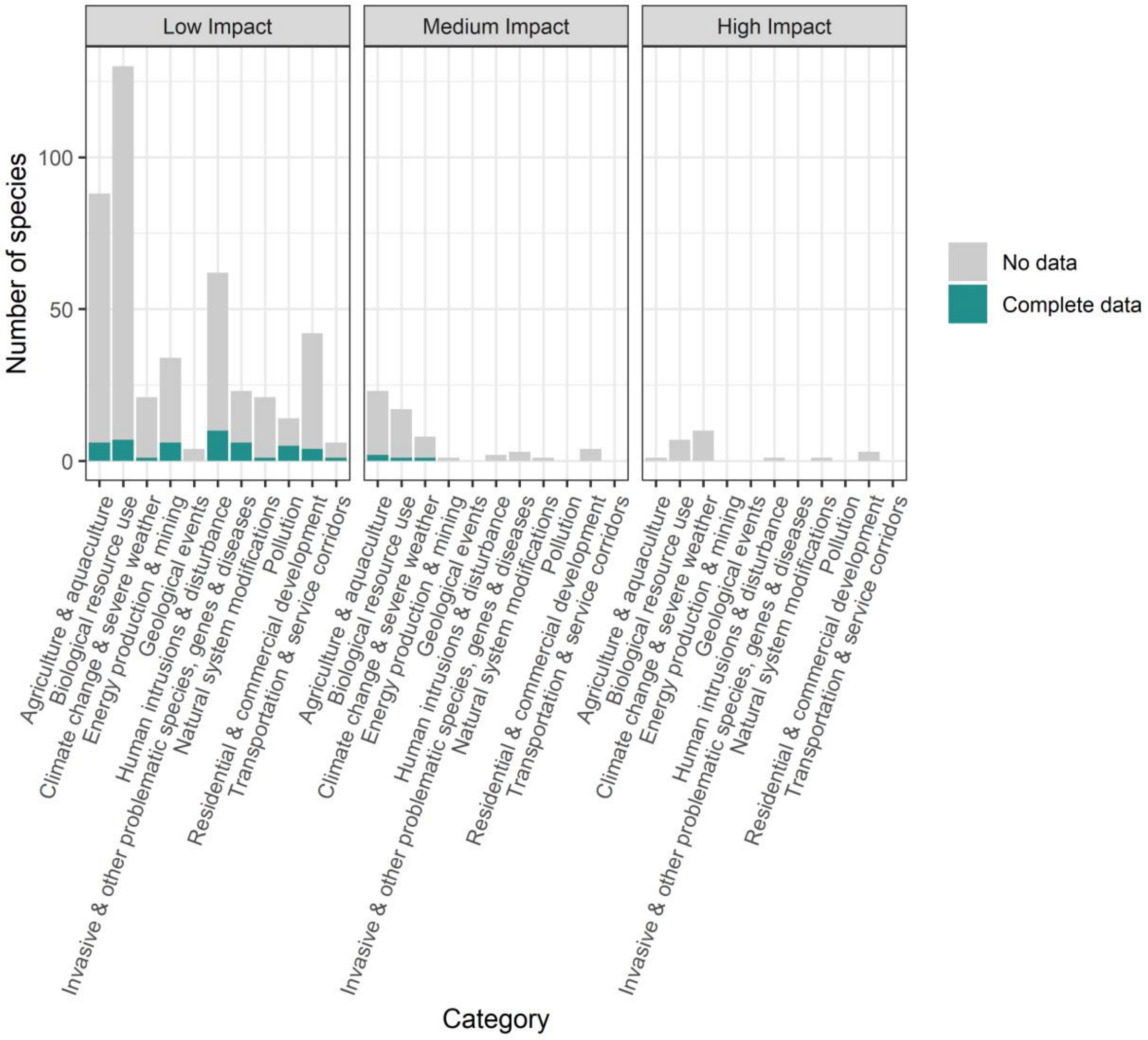


**Supplemental Figure S2** Bat species wing morphology data availability by IUCN threat type category and by assessed scale of the threat.

**Supplemental Table S3** Trait coverage for aspect ratio, wing loading, and relative wing loading by Ecoregion (WWF Ecoregions)

| **Trait** | **Ecoregion** | **Trait coverage** |
|---|---|---|
| Aspect ratio | Afrotropical | 28.50% |
| | Australasian | 23.10% |
| | Indomalayan | 31.80% |
| | Nearctic | 65.40% |
| | Neotropical | 28.30% |
| | Palearctic | 37.50% |
| Wing loading | Afrotropical | 29.20% |
| | Australasian | 23.10% |
| | Indomalayan | 31.10% |
| | Nearctic | 65.40% |
| | Neotropical | 28.10% |
| | Palearctic | 38.30% |
| Relative wing loading | Afrotropical | 28.20% |
| | Australasian | 23.10% |
| | Indomalayan | 31.80% |
| | Nearctic | 65.40% |
| | Neotropical | 29.10% |
| | Palearctic | 38.30% |
